# Modulation and Neural Correlates of Postmating Sleep Plasticity in *Drosophila* Females

**DOI:** 10.1101/2023.01.11.523670

**Authors:** José M. Duhart, Joseph R. Buchler, Sho Inami, Kyle J. Kennedy, B. Peter Jenny, Dinis J.S. Afonso, Kyunghee Koh

## Abstract

Sleep is essential, but animals may forgo sleep to engage in other critical behaviors, such as feeding and reproduction. Previous studies have shown that female flies show decreased sleep after mating, but our understanding of the process is limited. Here, we report that postmating nighttime sleep loss is modulated by diet and sleep deprivation, demonstrating a complex interaction among sleep, reproduction, and diet. We also report that female-specific pC1 neurons and sleep-promoting dorsal fan-shaped body (dFB) neurons are required for postmating sleep plasticity. Activating pC1 neurons leads to sleep suppression on standard fly culture media but has little sleep effect on sucrose-only food. Published connectome data suggest indirect, inhibitory connections among pC1 subtypes. Using calcium imaging, we show that activating the pC1e subtype inhibits dFB neurons. We propose that pC1 and dFB neurons integrate the mating status, food context, and sleep drive to modulate postmating sleep plasticity.

**Highlights:**

- Diet and sleep drive modulate female nighttime postmating sleep loss
- Female-specific pC1 neurons are required for postmating sleep loss
- Sleep-promoting dFB-projecting neurons are required for postmating sleep loss
- Activating pC1 subtypes promotes wakefulness and inhibits dFB-projecting neurons

**eTOC blurb:** Animals evaluate environmental conditions and internal states to make behavioral choices. Duhart et al. show that the decision to stay awake after mating in *Drosophila* females is modulated by food composition and sleep history and mediated by female-specific pC1 neurons acting upstream of the dFB sleep center.

## Introduction

While sleep serves essential functions, and its loss can lead to impaired fitness, animals may forgo sleep to engage in goal-directed behaviors critical for the survival of the individual or the species^1,2^. In *Drosophila*, male flies suppress sleep when mating opportunities arise^3–5^, and females suppress sleep after mating^6–9^. Since sleep loss can negatively impact health and cognition^10–13^, the balance between sleep and reproductive needs may be modulated by factors that affect the outcome of reproductive behavior. For instance, female-induced sleep suppression is attenuated when male flies have experienced a poor nutritional condition, which may indicate a low probability of offspring survival^14^. Additionally, when sleep pressure is increased by sleep deprivation, the presence of a female no longer leads to sleep loss^3^. These findings suggest that male flies integrate multiple sources of information to balance sleep and reproductive needs.

In addition to sleep suppression, mating leads to other behavioral changes in female flies, including increased egg laying and preference for protein-rich food^15^. These oviposition-related postmating changes can be modulated by the nutritional content and hardness of the egg-laying substrate, as well as by internal nutritional status^16–18^. These findings demonstrate that food composition confers relevant contextual information that affects how female flies behave after mating. Food containing sucrose only is sufficient for the survival of adult flies but not larvae, and mated females prefer not to lay eggs on sucrose-only substrates^16^. Thus, if oviposition-related activities contribute to postmating sleep loss, sleep may also be modulated by food type. Although a previous study found that food type does not influence postmating sleep loss^7^, the study examined only daytime sleep, and food type may modulate postmating sleep during the nighttime.

Recent studies have identified several neural substrates for integrating sleep, mating drives, and nutritional status in male flies. They include a subset of octopaminergic neurons, DN1p clock neurons, and a male-specific subset of Doublesex-positive pC1 cluster, P1 neurons^3–5^. Activation of P1 neurons or their postsynaptic partners, a pair of dopaminergic neurons projecting to the protocerebral bridge (DA-PB), leads to sleep reduction in normally-fed but not yeast-deprived males^14^, suggesting their role in nutritional modulation of the sleep-courtship balance. Additional PB-projecting neurons in the central complex regulate sleep in males^14^, adding to the growing number of central complex neurons involved in sleep regulation^19,20^.

The neural mechanisms conveying the mating status information from the peripheral sensory neurons to the central brain of female flies are well understood. Sex Peptide, which males deposit during copulation in the female ovary, inhibits the Sex Peptide sensory neurons (SPSN) and their postsynaptic partners in the abdominal ganglion (SAG)^21–24^. Recent studies have elucidated the circuits underlying select aspects of postmating behavioral changes, including feeding preferences, oviposition, and sexual receptivity^25–28^. Female-specific Doublesex-positive pC1 neurons, a counterpart to the male-specific P1 neurons, act downstream of the SPSN-SAG pathway to mediate postmating changes in oviposition and receptivity^25,26^. Previous work has shown that SPSN and SAG neurons are required for female postmating daytime sleep changes on a sucrose-only diet^29^. However, their involvement in nighttime sleep changes on a normal diet has not been examined. Furthermore, the role of pC1 neurons and previously described sleep-regulatory neurons in postmating sleep loss has yet to be investigated. The electron microscopy (EM)-based connectome of the adult female brain facilitates the investigation of the neural correlates of female-specific behavior^30,31^.

In the present work, we found that female flies exhibited substantial postmating sleep loss during the night on standard fly culture media (normal food) but not on sucrose-only food and that sleep deprivation attenuated postmating nighttime sleep suppression. We also found that SPSN, SAG, and female-specific pC1 neurons, as well as sleep-promoting dorsal fan-shaped body (dFB) neurons, are required for the effects of mating status on nighttime sleep. Activating female-specific pC1 neurons suppressed sleep on normal food but had little effect on sucrose-only food, suggesting that they play a role in the dietary modulation of postmating sleep plasticity. The available connectome of a female fly brain shows substantial inhibitory connections among pC1 subtypes via GABAergic OviIN neurons known for oviposition suppression^25,30^. We found that activating pC1e neurons, which receive inhibitory inputs from OviIN neurons, led to sleep suppression. Furthermore, calcium imaging demonstrated an inhibitory functional connection between the pC1e and dFB neurons. Our results show that postmating sleep changes are modulated by food context and sleep drive and suggest that information regarding the mating status, diet, and sleep history are integrated by pC1 and dFB neurons to regulate sleep in female flies.

## Results

### Food type modulates postmating sleep loss

To evaluate the effects of food type on postmating sleep suppression, we assessed sleep in virgin and mated female flies on two food substrates: normal laboratory food containing yeast, molasses, and cornmeal or 5% sucrose food. Virgin females raised on normal food were grouped with males to allow mating or kept in female-only groups for 24h before being individually placed in a glass tube containing normal food or 5% sucrose food (Figure 1A). We employed the multi-beam *Drosophila* Activity Monitor (multi-beam DAM) system with 17 infrared (IR) detectors previously used for assessing postmating sleep^32,33^. Mating led to reduced daytime sleep both in flies with access to 5% sucrose food and to normal food (Figures 1B–1D), as previously described^33^. The previous work did not examine the effects of food type on nighttime sleep. Interestingly, we found that mated females exhibited marked nighttime postmating sleep suppression on normal food but not on 5% sucrose food (Figures 1B–1E). These findings suggest that food composition modulates nighttime sleep specifically; thus, we focus our attention on nighttime sleep in subsequent experiments.

**Figure 1.**
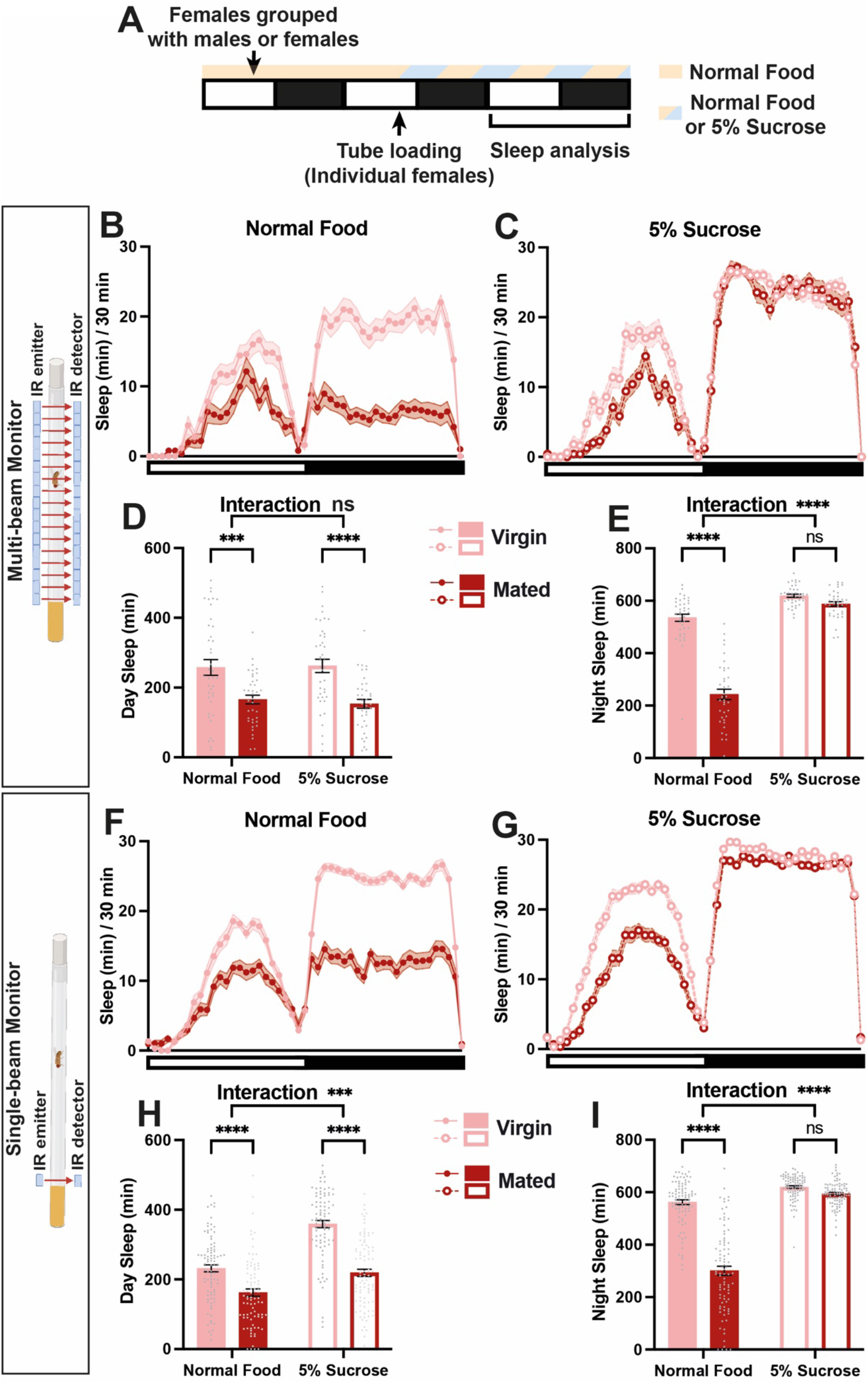
Food type modulates postmating nighttime sleep loss. **(A)** Experimental design. **(B** and **C)** Sleep profiles of virgin and mated females with access to normal food (B) or 5% sucrose (C), measured using multi-beam monitors. N = 38-40. **(D and E)** Daytime (D) and nighttime (E) sleep for flies shown in (B) and (C). **(F** and **G)** Sleep profiles of virgin and mated females with access to normal food (F) or 5% sucrose (G), measured using single-beam monitors. N = 88-91. **(H and I)** Daytime (H) and nighttime (I) sleep for flies shown in (F) and (G). Wild-type (Iso31) flies were used in all panels. In this and subsequent figures, bar graphs represent mean ± SEM, and gray symbols represent individual measurements. The white and black bars in the experimental design and below the x-axis in sleep profile graphs indicate 12-h light and 12-h dark periods, respectively. ***p<0.001, ****p<0.0001 and ns: not significant. Two-way ANOVA followed by Sidak’s post hoc test (D, E, H, and I). See also Figure S1.

We next tested whether the single-beam DAM system, suitable for high-throughput analysis, could also detect the effects of mating status and food composition on female sleep. Since mated females spend more time near food than virgin females^9^, we placed the IR sensor at a ~0.5 body distance from the food. We found that the single-beam data reproduced all the essential features of multi-beam data (Figures 1F–1I). Importantly, the single-beam data showed post-mating nighttime sleep loss on normal food but not on 5% sucrose food. Since single-beam monitors allow the analysis of more flies simultaneously than multi-beam monitors, we performed the remaining experiments using single-beam monitors unless otherwise stated.

Given that normal food but not sucrose-alone food allows for proper larval development, we wondered whether larval movement contributed to the apparent increase in wakefulness of mated females on normal food. To assay larval locomotion without adult movement, we removed adult females 24 h after placing them in monitor tubes (Figure S1A). We found that larval movement had a negligible contribution to sleep measurements during the time window used for sleep analysis in our experiments (Figures S1B–S1E). Moreover, analysis of nighttime sleep within 24 h of loading the flies into monitor tubes (larvae hatch ~24 h after eggs are laid), also showed that mated flies suppress their sleep on normal food but not on 5% sucrose (Figures S1F–S1H).

A recent study reported that a hidden Markov model could distinguish deep and light sleep states using DAM locomotor activity data^34^. We employed the model to test whether mated females slept less but more deeply than virgin females. We found that mated females on normal food exhibited a marked reduction in deep sleep at night without significantly altering the amount of light sleep (Figures S1H–S1K). This finding indicates that mated females do not compensate for short sleep by sleeping deeply. Together, our results show an interaction between food composition and mating status in regulating female sleep such that mated females show strong suppression of deep nighttime sleep when they have access to nutrient-rich food.

**Figure S1.**
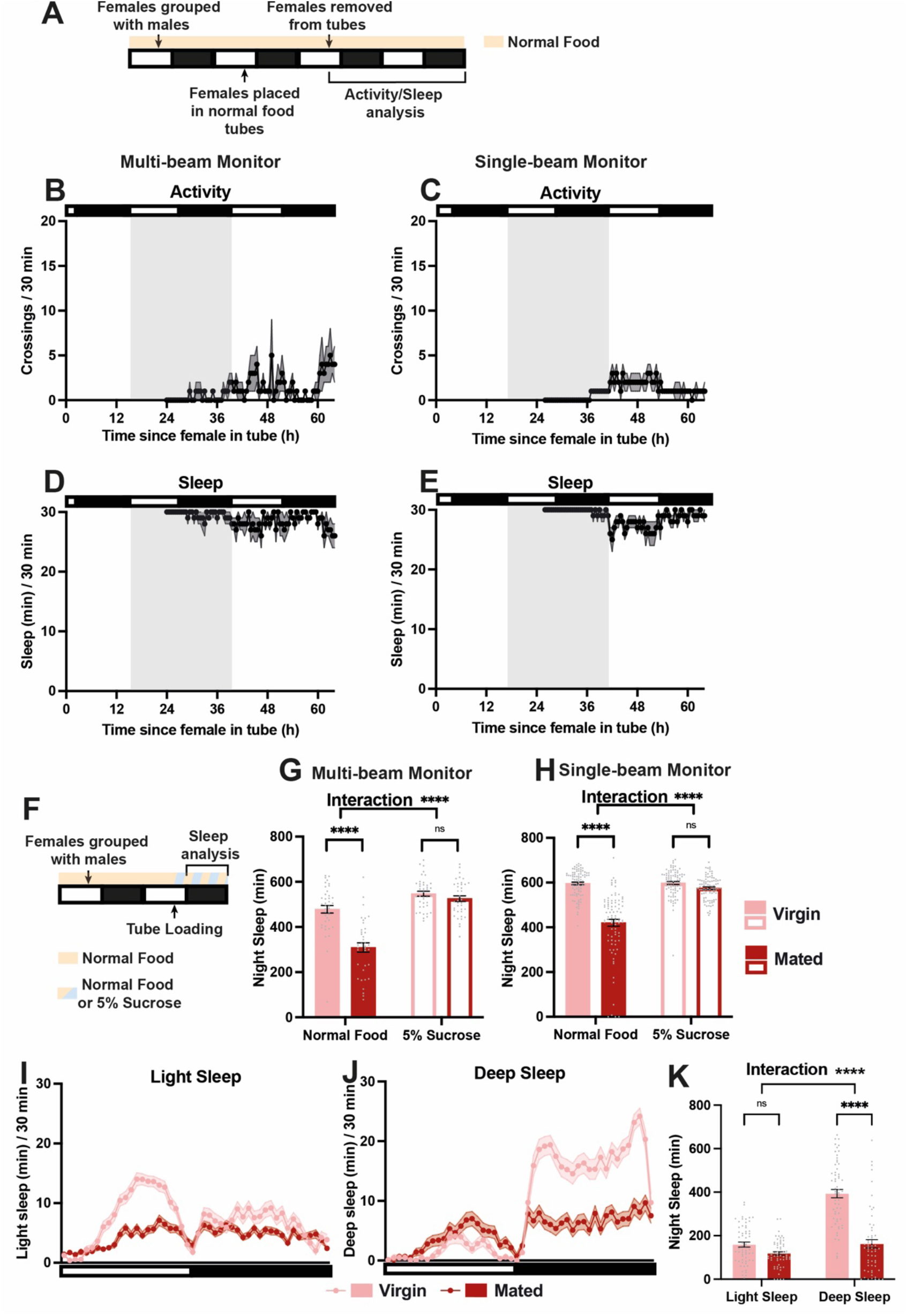
Postmating sleep suppression on normal food is not an artifact of larval movement and reflects changes in deep sleep. Related to Figure 1. **(A)** Experimental design for (B – E). **(B** and **C)** Activity measurements of hatched larvae using multi-beam (B) or single-beam (C) monitors. N = 22-24 for multi-beam monitors, and n = 88-91 for single-beam monitors. **(D** and **E)** Sleep measurements of hatched larvae using multi-beam (D) or single-beam (E) monitors shown in (B) and (C). **(F)** Experimental design for (G) and (H). **(G** and **H)** Nighttime sleep measured with multi-beam (G) and single-beam (H) monitors during the first night (ZT 12-24) after loading female flies shown in Figure 1 into glass tubes at ~ZT 8. **(I** and **J)** Sleep profiles for light (I) and deep (J) sleep of virgin and mated females on normal food shown in Figure 1F. ****p<0.0001 and ns: not significant. Two-way ANOVA followed by Sidak’s post hoc test (G and H).

### Food composition modulates postmating changes in egg laying and nighttime sleep

A proposed reason for reduced sleep in mated females is the need to lay eggs^9^. Female flies lay eggs on food substrates, and it is well documented that the number of eggs laid is strongly affected by the quality of the food substrate^16,35^. Indeed, analysis of our multi-beam DAM data showed that mated females spent more time next to normal food than virgin females, especially during the night (Figures 2A and S2). As noted above, mated females showed significantly reduced daytime sleep on 5% sucrose food (Figure 1; ^33^). In contrast, the time spent near 5% sucrose food does not differ between virgin and mated flies (Figure S2B – S2D). Activities close to food, such as egg laying and feeding, may be primarily responsible for nighttime postmating sleep loss on normal food. In contrast, during the daytime, on 5% sucrose, foraging for more nutritious food and better egg-laying substrates away from the food may contribute substantially to sleep loss.

**Figure 2.**
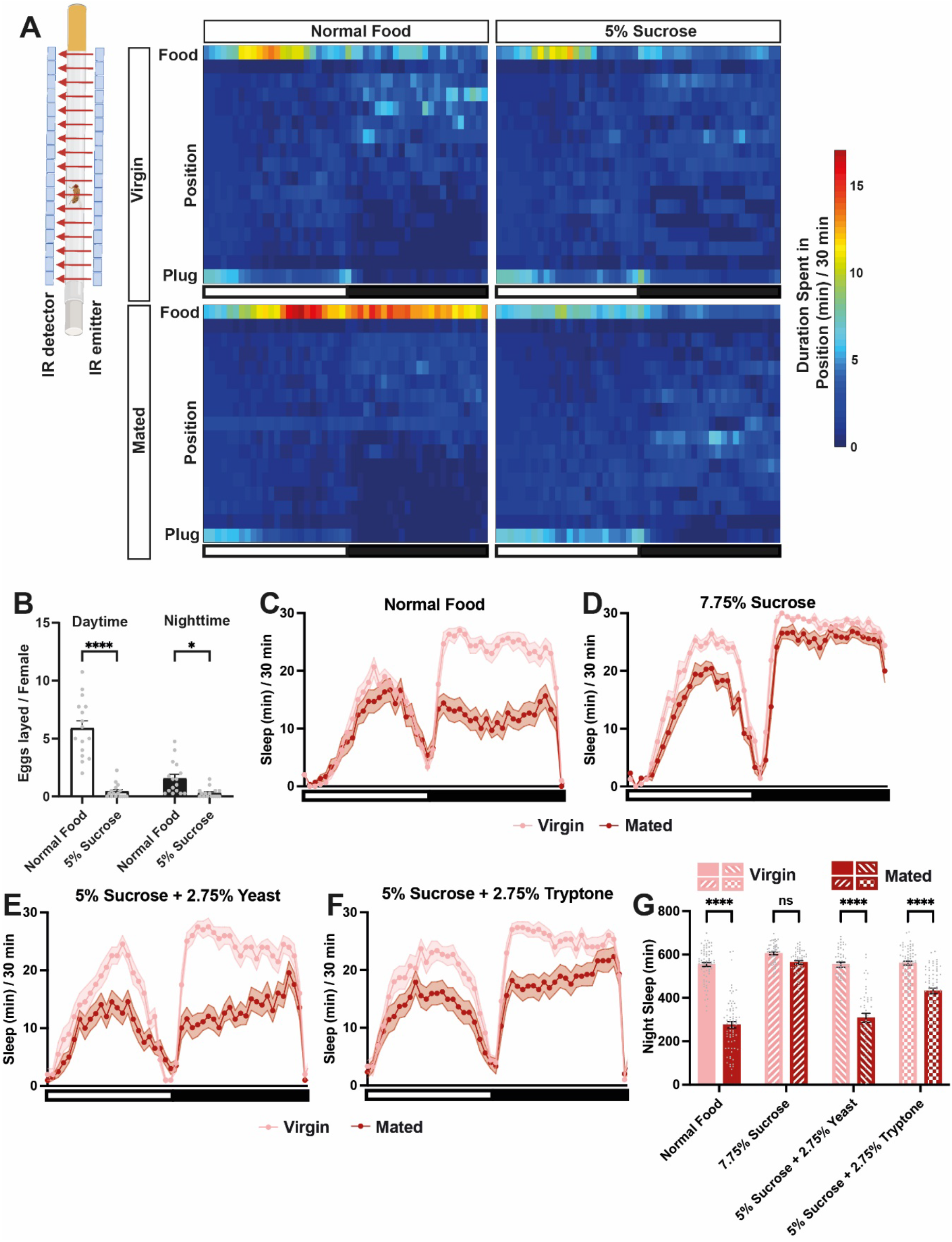
Food composition modulates postmating changes in nighttime sleep and egg laying. **(A)** Position heatmaps indicating the average duration that flies spent at different tube positions per 30-min bin using the multi-beam data shown in Figures 1B and 1C. The top of each heatmap (Food) is closest to the food. **(B)** Number of eggs laid per female on normal food or 5% sucrose. N = 16 vials (4 females/vial). **(C**-**F)** Sleep profiles of virgin and mated females with access to normal food (C), 7.75% sucrose (D), 5% sucrose + 2.75% yeast (E), or 5% sucrose + 2.75% tryptone (F). N = 47-72. **(G)** Nighttime sleep for flies in (C–F). Wild-type (Iso31) flies were used in all panels. *p<0.05, ****p<0.0001 and ns: not significant. Two-way ANOVA followed by Sidak’s post hoc test (B and G). See also Figure S2.

To examine whether nighttime sleep loss is associated with increased oviposition, we counted eggs laid by mated females during the daytime and nighttime on normal vs. sucrose food. We found that the number of eggs laid was significantly higher on normal food than on 5% sucrose during the nighttime as well as daytime (Figure 2B). This result suggests that oviposition-related activities account for at least some nighttime sleep loss after mating on normal food.

**Figure S2.**
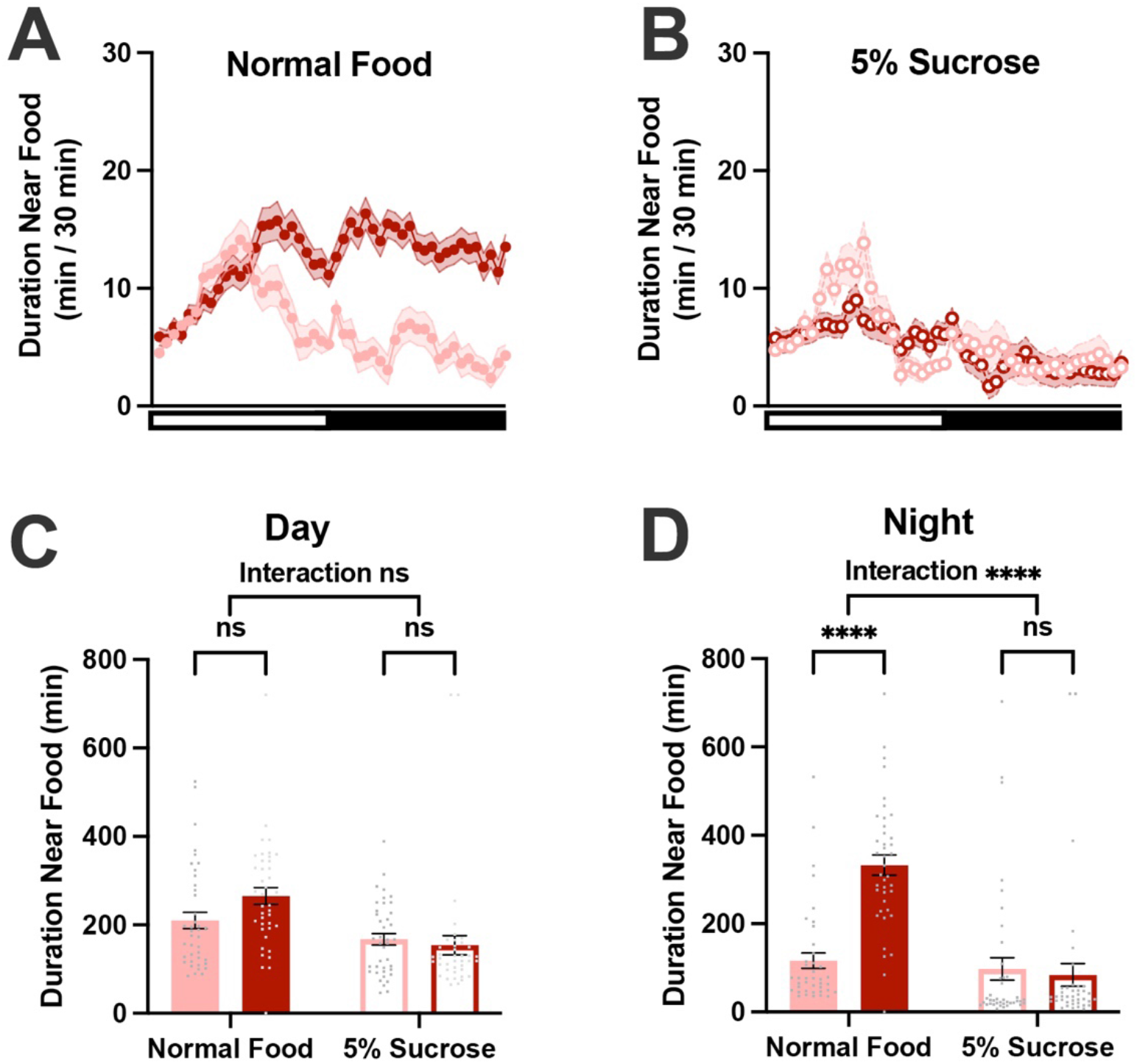
Mated females spend more time near normal food. Related to Figure 2. **(A and B)** Duration spent in the position closest to normal (A) or 5% sucrose (B) food per 30-min bin. Multi-beam data for flies shown in Figure 2A were used for the analysis. **(C and D)** Daytime (C) and nighttime (D) total duration spent near food for flies in (A and B). ****p < 0.0001 and ns: not significant. Two-way ANOVA followed by Sidak’s post hoc test.

Yeast is the primary source of protein in a normal laboratory fly diet and provides nutrients necessary for larval growth and survival^36^. In addition, mated females increase their preference for protein-rich food after mating^37,38^. Thus, we tested whether yeast and protein are nutritional components relevant for postmating nighttime sleep suppression. We found that supplementing 5% sucrose food with either yeast or tryptone (a mixture of peptides) was sufficient for strong postmating sleep suppression (Figures 2C and 2E–2G). This effect was not due to the altered caloric content of the diet since increasing the sucrose concentration to 7.75% to match the caloric content of the 5% sucrose + 2.75% tryptone diet did not lead to a significant effect of mating on nighttime sleep (Figures 2D and 2G). These findings point to yeast and protein as critical nutritional components for postmating sleep suppression.

### Sleep drive modulates the impact of mating on nighttime sleep

Sleep deprivation has previously been shown to attenuate female-induced sleep suppression in male flies^3^, demonstrating competition between sleep and sex drives in males. We wondered whether sleep deprivation would also attenuate postmating nighttime sleep loss in females, pointing to an analogous competition between sleep and reproductive drives in female flies. We subjected virgin and mated flies on normal food to 6 h of sleep deprivation at the beginning of the night via mechanical stimulation and examined their sleep amount during the remaining nighttime (Figure 3A). Comparable levels of sleep between virgin and mated flies were observed during the first hour after sleep deprivation (Figures 3B–3D). Although the mated females slept less than virgin females over the next several hours, the magnitude of the mating effect on nighttime sleep is reduced by sleep deprivation, as evidenced by the strong interaction between sleep deprivation and mating status (Figures 3B, 3C, and 3E). These results suggest sleep and reproductive drives compete such that unusually high sleep drive can override reproductive drive.

**Figure 3.**
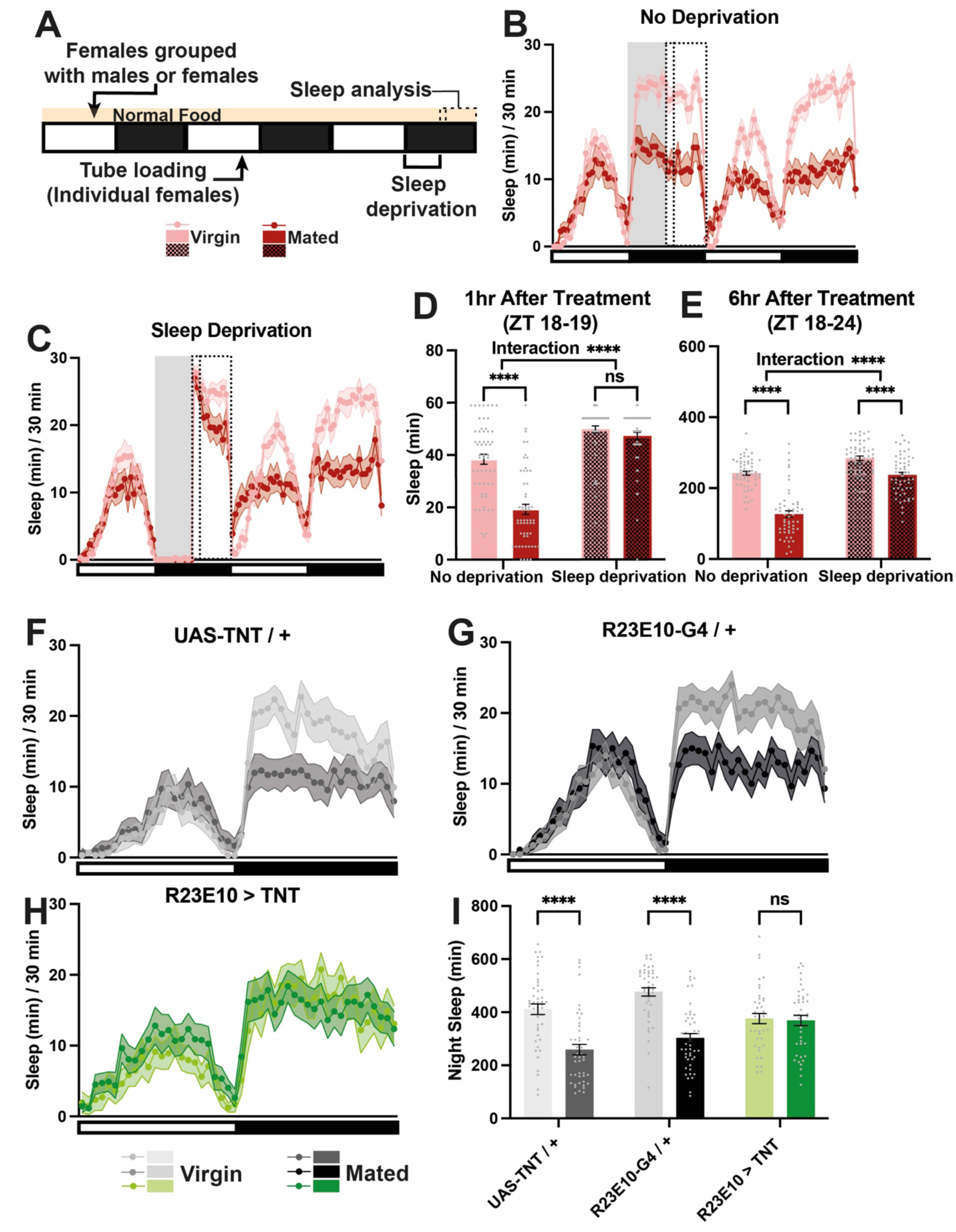
Sleep deprivation and the inhibition of dFB-projecting neurons attenuate the impact of mating on nighttime sleep. **(A)** Experimental design for panels (B–E). **(B**–**C)** Sleep profiles of Iso31 virgin and mated females on normal food, without (B) or with (C) mechanical sleep deprivation from ZT 12 to ZT 18 (gray rectangle). Dotted rectangles represent the 1-h and 6-h period after deprivation (ZT 18-19 and ZT 18-24). N = 55-56. **(D-E)** Sleep amounts during the 1-h (D) or 6-h (E) period after sleep deprivation for flies in (B) and (C). **(E**-**G)** Sleep profiles of virgin and mated females with impaired neurotransmitter release from dFB-projecting neurons (R23E10 > TNT, G) and their parental controls (E and F) on normal food. N = 41-48. **(H)** Nighttime sleep for flies in (E–G). ****p<0.0001 and ns: not significant. Two-way ANOVA followed by Sidak’s post hoc test (D, E, and H). See also Figure S3.

A group of neurons projecting to the dorsal fan-shaped body (dFB) play a critical role in the homeostatic response to sleep loss^39,40^. Since postmating sleep loss can be modulated by sleep deprivation, we tested the role of dFB neurons in postmating sleep regulation. We blocked neurotransmitter release from dFB neurons by expressing the tetanus toxin (TNT) light chain under the control of R23E10-Gal4^39^. This manipulation abolished the difference in nighttime sleep between virgin and mated females on normal food (Figures 3F–3I). Inhibiting dFB neurons had little effect on nighttime sleep in virgin and mated females on 5% sucrose (Figures S3A–S3D), a condition in which control females do not exhibit postmating nighttime sleep suppression. These results suggest that mating induces nighttime sleep suppression on normal food by inhibiting dFB neurons. We next examined the effects of activating dFB neurons by expressing the bacterial sodium channel NaChBac^41^ under the control of R23E10-Gal4. We found a greater sleep increase in mated females compared to virgin females (Figures S3E–S3G), as indicated by the highly significant interaction between genotype and mating status. These results suggest that increasing sleep pressure, either by sleep deprivation or through dFB neuronal activation, attenuates the sleep-suppressive effect of mating in females with access to normal food.

**Figure S3.**
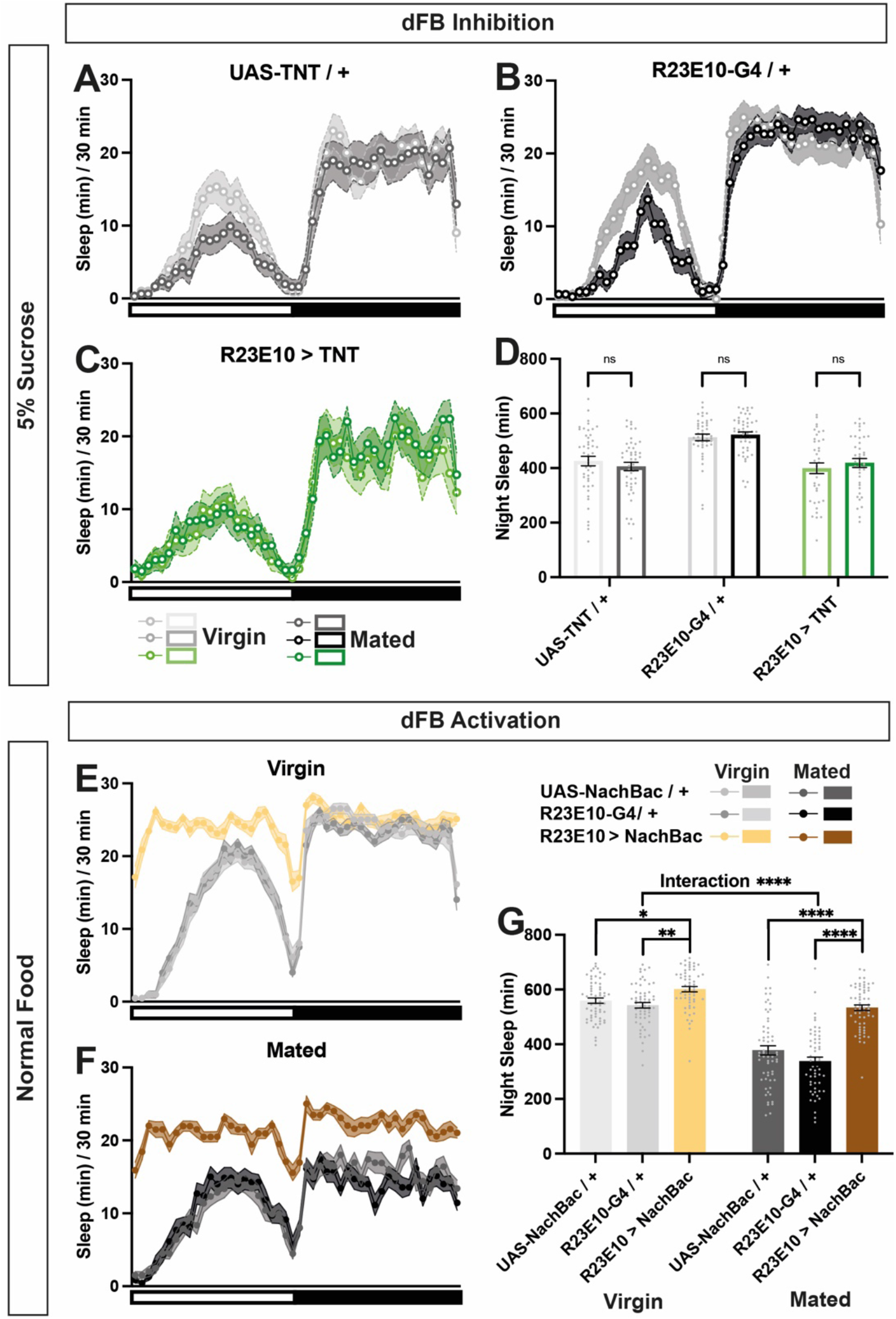
dFB-projecting neurons modulate postmating sleep loss. Related to Figure 3. **(A**–**C)** Sleep profiles of virgin and mated females with impaired neurotransmitter release from dFB neurons (C) or their parental controls on 5% sucrose (A and B). N = 38-48. **(D)** Nighttime sleep for flies in (A–C). **(E** and **F)** Sleep profiles of virgin **(E)** or mated **(F)** females expressing the bacterial sodium channel NachBac in dFB neurons (R23E10 >NachBac) and parental controls (UAS-NachBac/+ and R23E10-G4/+) on normal food. N = 56-62. **(G**) Nighttime sleep for flies in (E) and (F). *p<0.05, **p<0.01, ****p<0.0001 and ns: not significant. Two-way ANOVA followed by Sidak’s post hoc test (D and G).

### pC1 neurons are required for postmating sleep suppression

Since mated flies on normal food show both increased egg laying and reduced sleep compared to those on 5% sucrose, we tested whether components of the oviposition circuit are also involved in postmating nighttime sleep suppression^23,25,27,42^. The neural substrates acting downstream of SPSN and SAG neurons in the central brain for oviposition include female-specific pC1, OviIN, OviEN, and OviDN neurons (Figure 4A; ^25^). As mentioned, SPSN and SAG neurons, the first two neuronal groups in this circuit, mediate daytime postmating sleep suppression on 5% sucrose food^43^. We found that impairing their neurotransmitter release prevented postmating nighttime sleep changes in female flies on normal food (Figure S4), indicating that they also mediate nighttime sleep changes.

**Figure 4.**
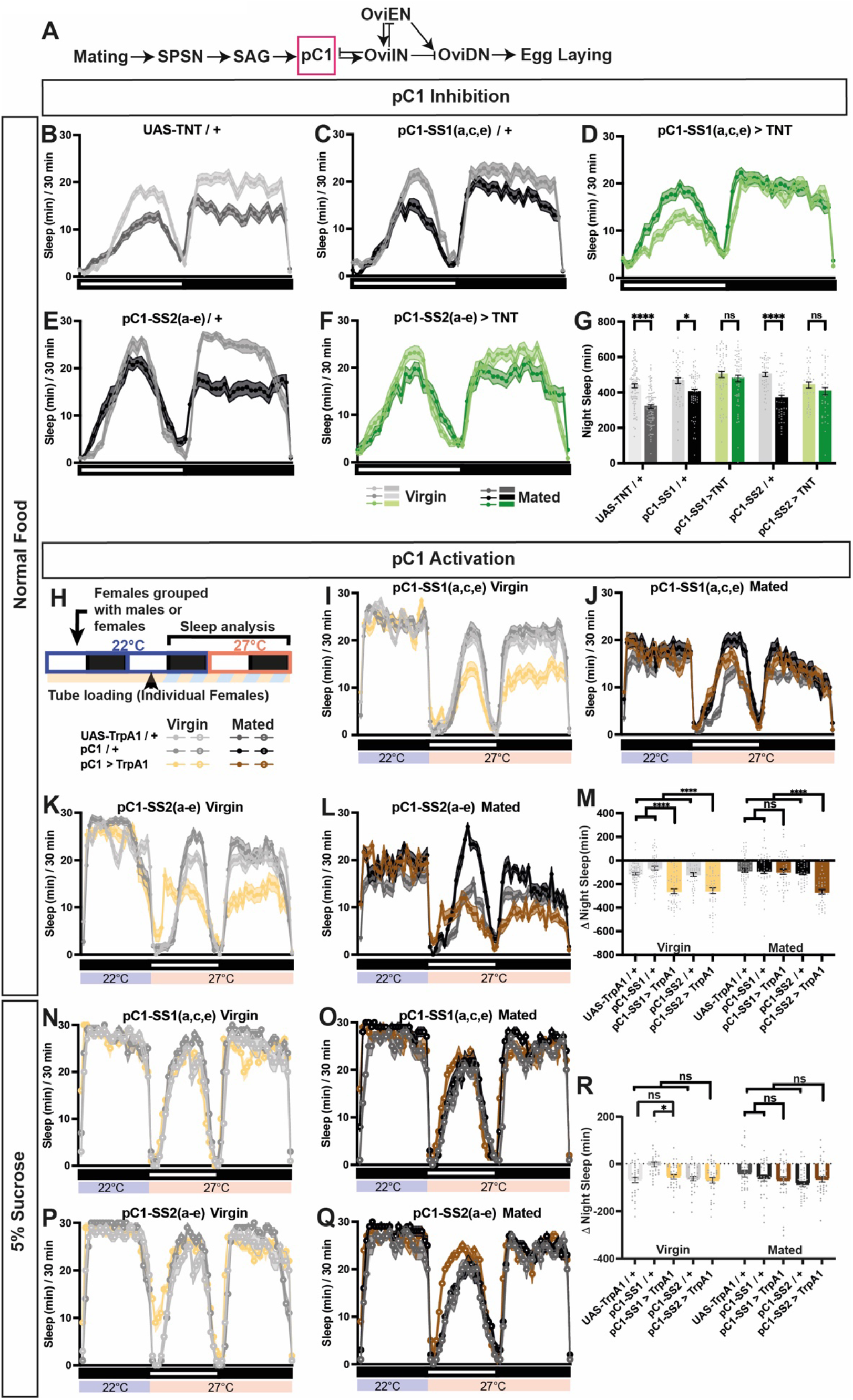
pC1 neurons are required for postmating sleep suppression. **(A)** Neural circuit recruited for increased egg laying after mating from ^25^. **(B**–**F)** Sleep profiles of virgin and mated females with impaired neurotransmitter release from pC1a,c,e subtypes (pC1-SS1(a,c,e) >TNT, D) or all pC1 subtypes (pC1-SS2(a-e) > TNT, F) and their parental controls (B, C, and E) on normal food. Split Gal4 lines, pC1-SS1 and pC1-SS2 ^25^, were used to target pC1a,c,e, and pC1a-e, respectively. N = 41-95. **(G)** Nighttime sleep for flies in (B–F). **(H)** Experimental design for (I–R). (**I–L**) Sleep profiles of virgin (I and K) or mated (J and L) females expressing the warmth-activated TrpA1 channel in pC1 (pC1-SS1 (a,c,e) > TrpA1 or pC1-SS2(a-e) > TrpA1) neurons and their parental controls on normal food. TrpA1 was activated by raising the temperature from 22°C to 27°C. N = 42–64. **(M)** Nighttime sleep change (sleep at 27°C – baseline sleep at 22°C) for flies in (I–L). **(N–Q)** Sleep profiles of virgin (N and P) or mated (O and Q) females of the same genotypes as (I–L) on 5% sucrose. N = 30-32. **(R)** Nighttime sleep change (sleep at 27°C – baseline sleep at 22°C) for flies in (N–Q). *p<0.05, ****p<0.0001 and ns: not significant. Two-way ANOVA followed by Sidak’s (G) or Tukey’s (M, and R) post hoc test. See also Figure S4 and S5.

**Figure S4.**
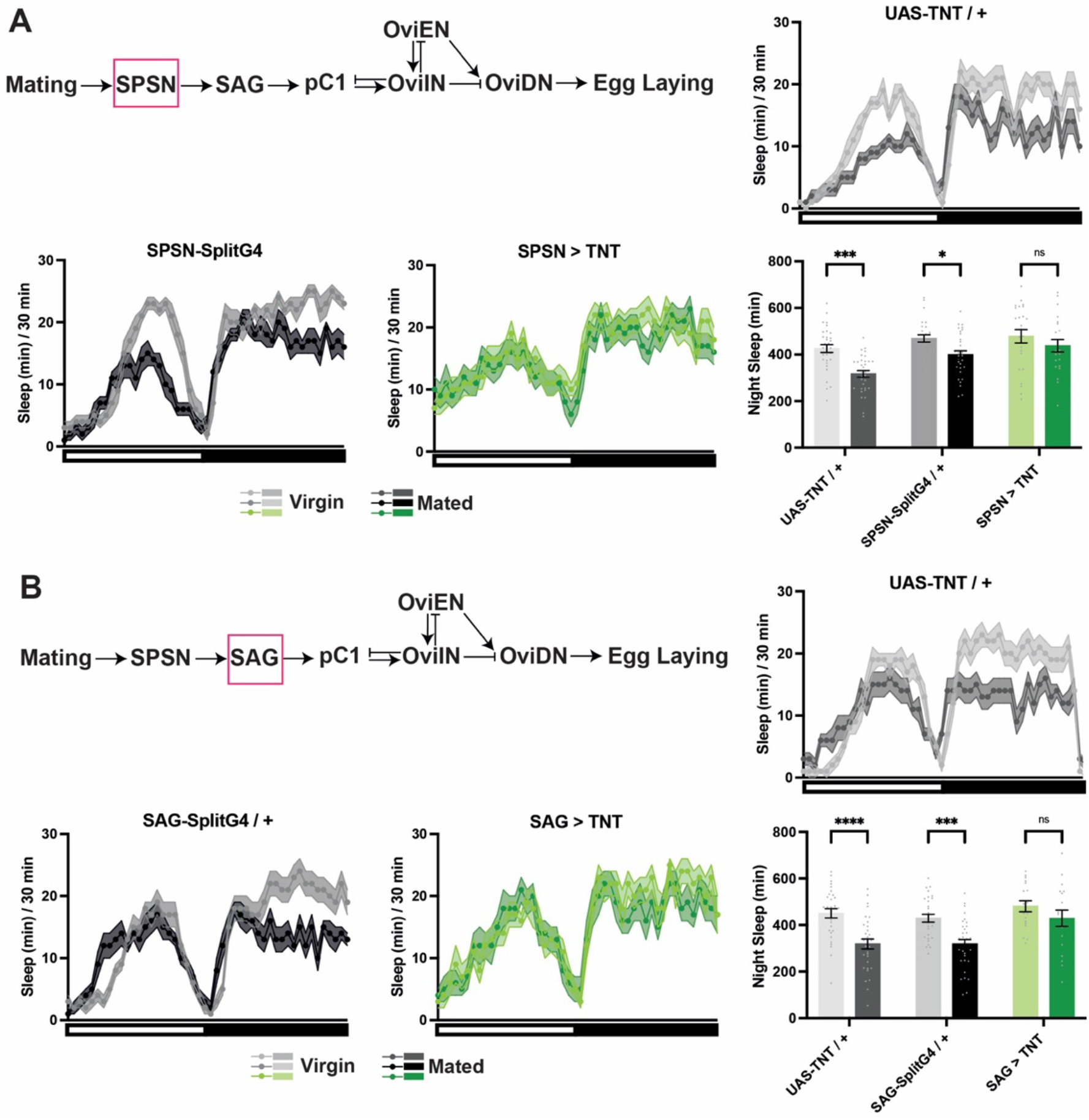
SAG and SPSN neurons are required for postmating sleep suppression. Related to Figure 4. **(A)** Sleep profiles and nighttime sleep of virgin and mated females with impaired neurotransmitter release from SPSN neurons (SPSN > TNT) and parental controls (UAS-TNT / + and SPSN-splitG4 / +) on normal food. N = 22-32. **(B)** Sleep profiles and nighttime sleep of virgin and mated females with impaired neurotransmitter release from SAG neurons (SAG > TNT) and parental controls (UAS-TNT / + and SAG-splitG4 / +) on normal food. N = 18-32. *p<0.05, ***p<0.001, ****p<0.0001 and ns: not significant. Two-way ANOVA followed by Sidak’s post hoc test.

Female-specific pC1 neurons receive mating status information from SAG neurons and control egg laying by acting on GABAergic OviIN neurons that inhibit egg laying (Figure 4A;^25^). OviIN neurons also receive information about the egg-laying substrate, including sugar content and hardness, and interact with OviEN neurons to control oviposition through the descending OviDN neurons^25^. There are five female-specific pC1 subtypes: a-e. We tested the effect of silencing pC1 neurons using two intersectional split-Gal4 drivers: pC1-SS1, which includes pC1 subtypes a, c, and e, and pC1-SS2, which includes all five pC1 cell subtypes^25^. We found that preventing neurotransmitter release from pC1 neurons using both pC1-SS drivers attenuated the difference in nighttime sleep between mated and virgin females (Figures 4B–4G). This finding indicates that signaling from pC1a, c, and e subtypes is necessary for postmating nighttime sleep changes.

We next activated pC1 neurons by expressing the warmth-sensitive cation channel TrpA1^44^ and elevating the temperature (Figure 4H). When flies had access to normal food, activation of pC1 subtypes a, c, and e suppressed sleep in virgin but not mated females (Figures 4I, 4J, and 4M). In contrast, activation of all five pC1 subtypes led to significant sleep reduction in both virgins and mated females (Figures 4K–4M), suggesting subtype-specific sleep-regulatory roles. Interestingly, in virgin and mated females with access to 5% sucrose, pC1 activation did not elicit significant changes in sleep (Figures 4N–4R), suggesting that pC1 neuron activity suppresses sleep only in specific food contexts. Silencing of neurons downstream of pC1 in the oviposition circuit, i.e., OviIN, OviEN, and OviDN neurons, did not prevent nighttime sleep suppression in mated flies (Figure S5). These results suggest that the Sex Peptide signal elicited by mating is conveyed through the SPSN, SAG, and pC1 neurons to influence sleep in a food composition-dependent manner.

**Figure S5.**
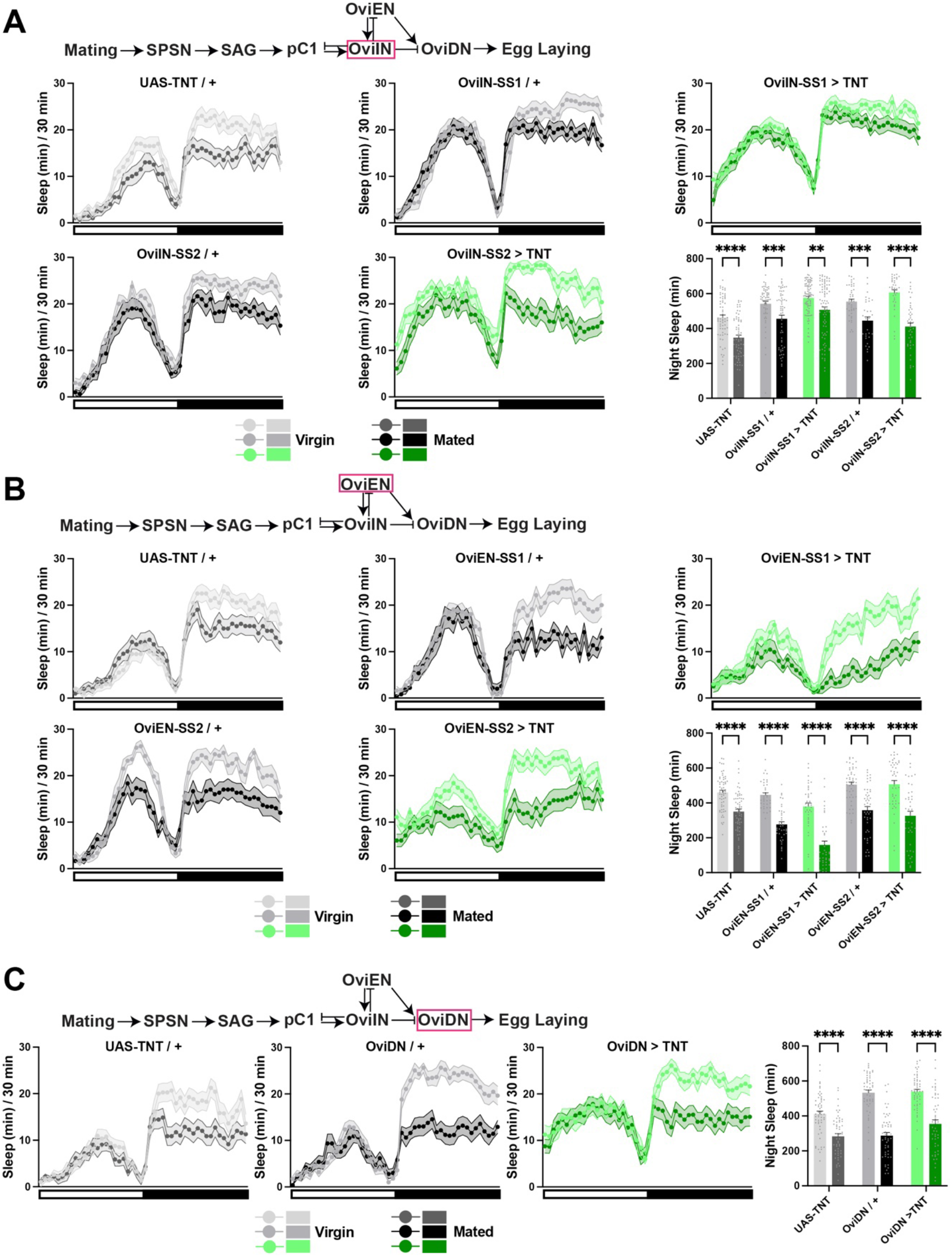
OviIN, OviEN and OviDN neurons are dispensable for postmating sleep suppression. Related to Figure 4. **(A)** Sleep profiles and nighttime sleep of virgin and mated females with impaired neurotransmitter release from OviIN (OviIN-SS1/2 > TNT) neurons and parental controls (UAS-TNT / +, OviIN-SS1 / +, and OviIN-SS2 / +) on normal food. N = 32-86. **(B)** Sleep profiles and nighttime sleep of virgin and mated females with impaired neurotransmitter release from OviEN (OviEN-SS12 > TNT) neurons and parental controls (UAS-TNT / +, OviEN-SS1 / +, and OviEN-SS2 / +) on normal food. N = 37-63. **(C)** Sleep profiles and nighttime sleep of virgin or mated females with impaired neurotransmitter release from OviDN (OviDN > TNT) neurons and parental controls (UAS-TNT / +, OviDN-SS2 / +) on normal food. N = 51-55. **p<0.001, ***p<0.001 and ****p<0.0001. Two-way ANOVA followed by Sidak’s post hoc test.

### pC1 subtypes have distinct roles in postmating sleep regulation

Activating pC1 neurons using pC1-SS1 and -SS2 suppresses egg laying in mated females^25^. Since egg laying is associated with wakefulness, we expected that pC1 activation would also suppress wakefulness. Unexpectedly, activating pC1a,c,e (using pC1-SS1) or pC1a-e (using pC1-SS2) led to sleep suppression (Figures 4H–4M), which could be explained by inhibitory connections among pC1 subtypes. Since all five female-specific pC1 subtypes are cholinergic, any inhibitory connections among pC1 subtypes may be indirect. pC1 neurons exhibit heterogeneity in their connections with SAG neurons, other pC1 subtypes, and GABAergic OviIN neurons^25^. Examination of synaptic connectivity data for pC1 subtypes and inhibitory OviIN neurons^25^ revealed that OviIN neurons provide many synaptic inputs to pC1c and pC1e. Our pC1 inhibition data (Figures 4B–4G) suggest that the a, c, and e subtypes included in both SS1 and SS2 drivers are required for sleep regulation by mating status. Previous work showed that stimulating SAG neurons had the most potent effect on pC1a activity and little effect on pC1c,e^25^. Thus, we looked for inhibitory connections between pC1a and pC1c,e via OviIN, and noted that pC1d is a good candidate for linking pC1a to OviIN and pC1c,e neurons. The connectome data and our findings suggest a model in which Sex Peptide sequentially inhibits SPSN, SAG, pC1a, pC1d, and OviIN neurons, which then disinhibits wake-promoting pC1c and pC1e neurons, resulting in postmating sleep loss (Figure 5A). However, given that OviIN neurons are dispensable for the mating effects on sleep (Figure S5A), an OviIN-independent pathway may also convey mating status information to sleep-regulatory neurons.

**Figure 5.**
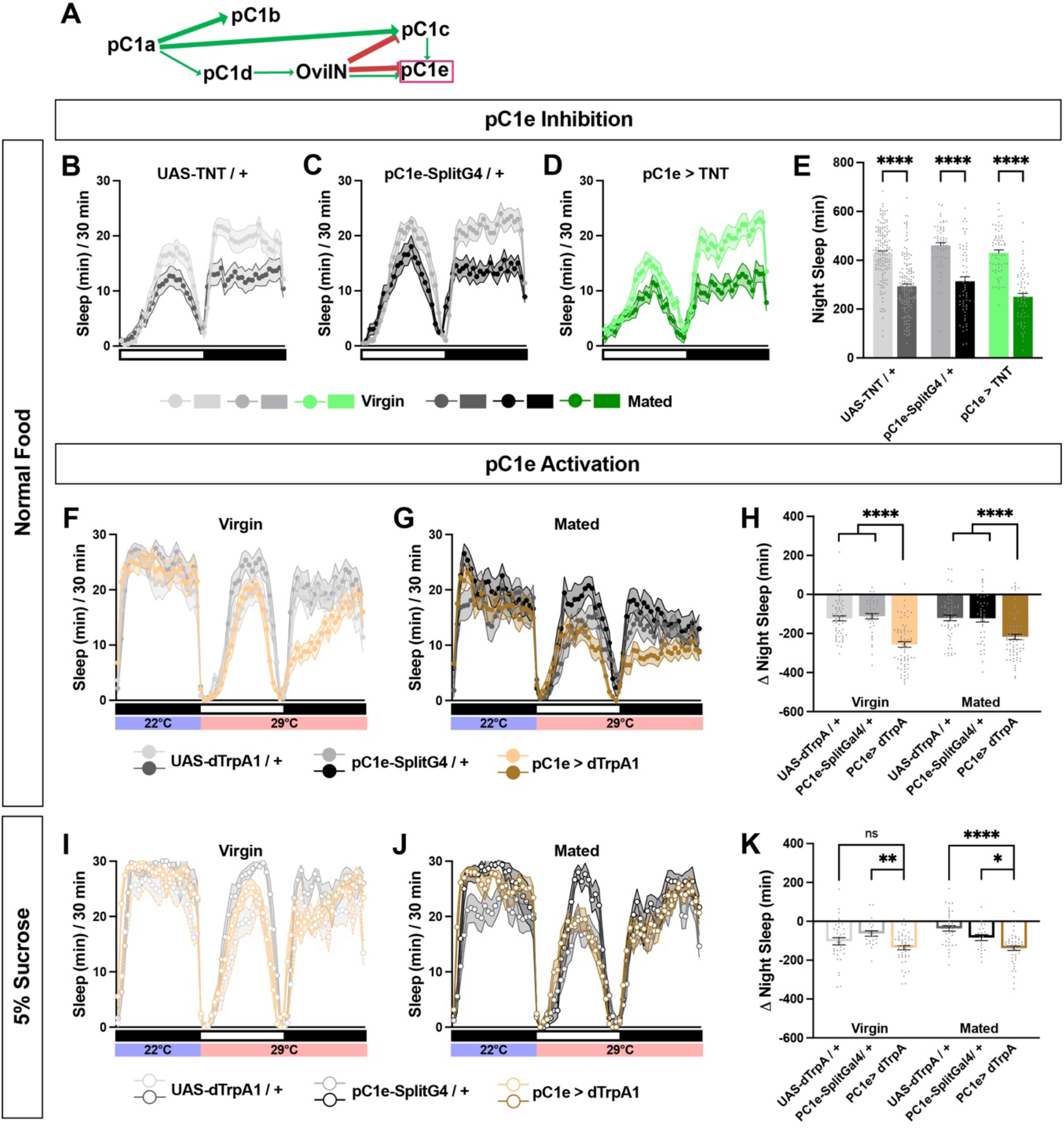
Activation of the pC1e subtype suppresses sleep. **(A)** Simplified diagram of the connections among pC1 and OviIN neurons based on an EM dataset (Wang et al., 2020). Only connections with over 10 synapses are shown. **(B–D)** Sleep profiles of virgin and mated females with impaired neurotransmitter release from pC1e (pC1e > TNT, D) and their parental controls (B and C) on normal food. N = 62-150. **(E)** Nighttime sleep change (sleep at 29°C – baseline sleep at 22°C) for flies in (B-D). **(F–G)** Sleep profiles of virgins (F) and mated (G) females expressing the warmth-activated TrpA1 channel in pC1e (pC1e > TrpA1) and their parental controls on normal food. TrpA1 was activated by raising the temperature from 22°C to 29°C. N = 42-73. **(H)** Nighttime sleep change (sleep at 29°C – baseline sleep at 22°C) for flies in (F) and (G). **(I–J)** Sleep profiles of virgins (I) and mated (J) females of the same genotypes as (F) and (G) on 5% sucrose food. TrpA1 was activated by raising the temperature from 22°C to 29°C. N = 21-49. **(K)** Nighttime sleep change (sleep at 29°C – baseline sleep at 22°C) for flies in (I) and (J). *p<0.05, **<0.01, ****p<0.0001 and ns: not significant. Two-way ANOVA followed by Sidak’s (E) or Dunnett’s (H and K) post hoc test.

To examine the role of pC1 subtypes in postmating sleep regulation, we employed a pC1e-specific split-gal4 driver^45^. (Due to a lack of specific drivers, we could not test the specific role of pC1a or pC1c.) Inhibiting pC1e neurons had little effect on postmating sleep changes (Figures 5B–5E), perhaps due to redundancy with pC1c. On the other hand, activating the pC1e subtype using TrpA1 expression and a warm temperature led to substantial sleep suppression in virgin and mated females on normal food (Figures 5F–5H). In contrast, activating pC1e had little effect on sleep in virgin females on 5% sucrose food (Figures 5I and 5K). Although activating pC1e in mated females resulted in a statistically significant decrease in sleep on sucrose food (Figures 5J and 5K), the magnitude of the sleep reduction is modest compared to that seen on normal food. These results suggest that pC1e activation suppresses sleep in a diet-dependent manner.

### dFB-projecting neurons act downstream of pC1 neurons for postmating sleep regulation

Given the requirement of both pC1 and dFB neurons for postmating nighttime sleep plasticity, we wondered whether the two groups of neurons are connected. The connectome data did not reveal any direct synaptic connection between the two groups. To test whether there are indirect functional connections, we expressed the ATP-sensitive P2X2 receptor in pC1a,c,e using the pC1-SS1 driver and the calcium sensor GCaMP7b in dFB neurons using the R23E10 driver, respectively. We dissected the brains and examined the GCaMP7b signal in the cell bodies of dFB neurons. Activating pC1a,c,e neurons with ATP perfusion led to significantly reduced calcium levels in dFB neurons (Figures 6A–6C). Data from virgin and mated females were pooled because we did not observe any effect of mating on the magnitude of dFB responses to pC1a,c,e activation. The lack of difference between virgin and mated females may be a consequence of using dissected brains, which lack Sex Peptide signaling from SPSN neurons in the ovary.

**Figure 6.**
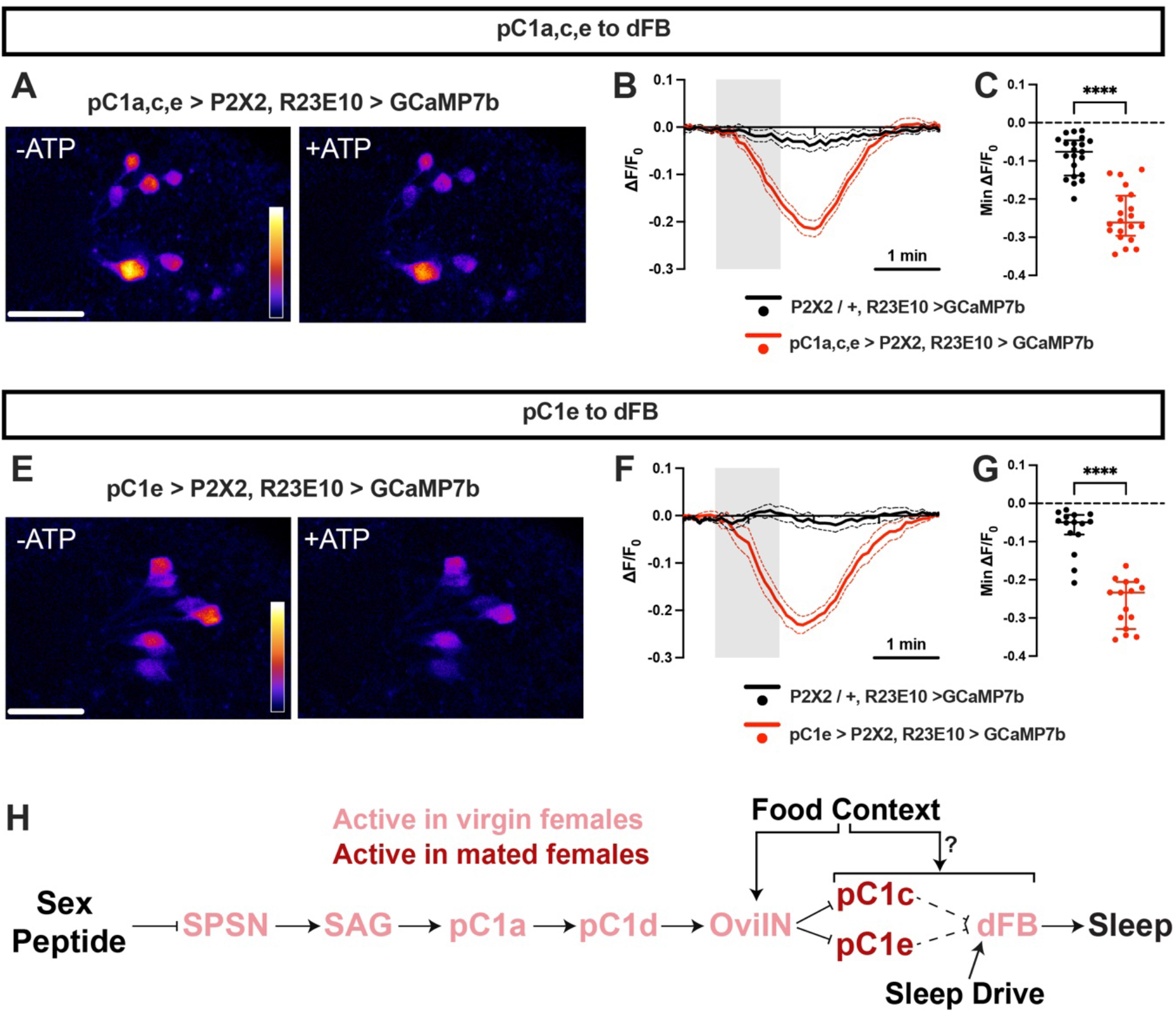
dFB-projecting neurons are inhibited by the activation of pC1a,c,e or pC1e neurons. **(A)** GCaMP7b signal in dFB-projecting neurons (R23E10 > GCaMP7b) before and shortly after ATP perfusion in a dissected female brain expressing the ATP-sensitive P2X2 receptor in pC1a,c,e neurons. The “fire” lookup table is used. Scale bar: 30 μm. **(B)** Fluorescence traces of normalized GCaMP7b signal (ΔF/F) in the cell bodies of dFB neurons in response to ATP perfusion in females expressing P2X2 in pC1a,c,e neurons (pC1a,c,e > P2X2). Flies carrying R23E10-LexA, LexAop-GCaMP7b, and UAS-P2X2, but not pC1-SS1(a,c,e), served as negative controls. N = 20-21 brains. **(C)** Minimum (Min) ΔF/F responses for brains in (B). **(D)** GCaMP7b signal in dFB-projecting neurons (R23E10 > GCaMP7b) before and shortly after ATP perfusion in a dissected female brain expressing P2X2 in pC1e neurons. The “fire” lookup table is used. Scale bar: 30 μm. **(E)** Fluorescence traces of normalized GCaMP7b signal (ΔF/F) in the cell bodies of dFB neurons in response to ATP perfusion in females expressing P2X2 in pC1e neurons (pC1e > P2X2). Flies carrying R23E10-LexA, LexAop-GCaMP7b, and UAS-P2X2, but not pC1e-splitG4, served as negative controls. N = 15 brains. **(F)** Minimum (Min) ΔF/F responses for brains in (E). **(G)** Model for postmating sleep plasticity. Sex Peptide deposited by males during copulation leads to the inhibition of SPSN, SAG, pC1a, pC1d, and OviIN. The inhibition of GABAergic OviIN neurons results in the disinhibition (activation) of pC1c,e subtypes. Activating pC1c,e neurons in mated females leads to the inhibition of the sleep-promoting dFB neurons through indirect connections (dotted lines). OviIN receives information about the sugar content ^25^. Information about additional nutrients may be integrated at the level of pC1c,e or their downstream neurons. We propose that dFB-projecting neurons integrate information regarding the mating status, food content, and sleep drive^39^ to regulate female sleep. ****p<0.0001. Unpaired Student’s t test (C, F).

We next employed a pC1e-specific split-Gal4 driver to examine the functional connectivity between pC1e and dFB neurons. We found that activating the pC1e subtype alone reduced calcium levels in dFB neurons (Figures 6D–6F), similar to activating pC1a,c,e together. The neural activity of the pC1e subtype may dominate when it is activated together with its upstream inhibitor, the pC1a subtype. The inhibitory connection between pC1e and dFB neurons is likely indirect since pC1 neurons are excitatory^25^, and pC1e activation leads to sleep suppression, while dFB activation promotes sleep^46^.

## Discussion

The circadian clock and sleep history are two primary regulators of sleep^47^. However, various factors, including food availability and sexual partners, modulate sleep^1,2^. For example, male arctic sandpipers greatly reduce their sleep time during a 3-week mating season^48^. Male flies also reduce sleep in the presence of female flies^3–5^. However, when male flies are deprived of yeast or protein, they no longer suppress sleep when female flies are around ^14^. These results suggest that animals evaluate the environmental context and internal states to determine if the benefits outweigh the costs of sleep loss. Since fly larvae cannot survive without protein, successful mating may not yield surviving progeny when protein is scarce, reducing the benefits of courtship. Our results show that a tradeoff between sleep and reproductive activities in female flies is also modulated by food context. We found that female flies spent more time next to normal food upon mating, consistent with previous findings^9^. We did not observe a similar change in females on sucrose-only food, and thus the increased time near food may be related to two other postmating changes: increased egg laying and feeding preference for protein-rich food^15^. Sucrose-only food may decrease the drive for egg laying and feeding, reducing the need to sacrifice sleep in mated flies when food lacks proper nutrients for larval growth. Our work provides novel insights into the mechanisms underlying complex interactions among multiple factors influencing sleep.

Several studies have shown that sleep loss adversely affects cognition and survival^10–13^. Thus, sleep loss for goal-directed behaviors may be modulated to prevent excessive sleep debt. Indeed, we found that several hours of forced sleep deprivation led mated female flies to exhibit rebound sleep over the next few hours. On the other hand, female flies suppressed daytime sleep for 6-8 days after mating^33,49^. In addition, our sleep depth analysis indicated that mated females did not trade long, light sleep with short, deep sleep. Together, these results suggest an altered homeostatic set point in mated females. Previous work revealed similar interactions between sleep and sex drives in male flies. Male sleep loss induced by female partners is attenuated by sleep deprivation^3^, and female pheromones suppress sleep in males so that they can maintain reduced sleep over multiple days^50^. A recent study found that most sleep in flies is not necessary for survival^9^. The optimal sleep amount may be context dependent. For example, the study of male arctic sandpipers mentioned above found that those who slept the least had the most progeny, indicating that sleep loss during mating periods provides a competitive advantage. In another example, whereas memory consolidation after olfactory conditioning requires increased sleep in well-fed flies, starved flies do not show elevated sleep after the same training without suffering from impaired memory^51^. Together, these findings suggest that significant changes in life circumstances, such as mating opportunities, copulations, and starvation, may result in homeostatic set point resetting.

Mating induces a variety of behavioral changes in female flies, including increased egg laying, preference for protein-rich food, and aggressiveness, as well as decreased sexual receptivity^15^. These changes depend on Sex Peptide, which is transferred from the males during copulation and acts on the SPSN-SAG circuit^23,25,27,28,42^. Previous work has shown that daytime female postmating sleep suppression on 5% sucrose also depends on SPSN and SAG neurons^7^. Our work shows that postmating nighttime sleep suppression on normal food also relies on SPSN and SAG neurons, as well as their downstream synaptic partners, female-specific pC1 neurons. Since Sex Peptide inhibits SPSN and SAG neurons, and SAG to pC1 connections are excitatory, it was unexpected that pC1 activation (associated with the virgin status) led to sleep suppression (associated with the mated status). However, since GABAergic OviIN neurons provide numerous inputs to pC1c and pC1e subtypes^25^, mating could lead to activation of pC1c,e through inhibition of OviIN neurons (Figure 6D). Consistent with the model, activating only pC1e neurons led to sleep suppression. Our model predicts that activating pC1c alone would also decrease sleep and that activating pC1a alone would increase sleep. However, testing them would require specific drivers for pC1c and pC1a. Our finding that inhibiting OviIN alone did not eliminate the effects of mating on sleep suggests that there may be additional pathways conveying mating information from SAG neurons to the dFB sleep center, both of which are required for postmating sleep loss.

Our examination of published connectome data^25^ revealed indirect inhibitory connections among pC1 subtypes. Consistent with the inhibitory connections, a previous study found that activating a subset of pC1 neurons had an inhibitory effect on female chasing behavior elicited by pC1d,e subtypes^52^. We found that simultaneously activating pC1a,c,e subtypes using pC1-SS1 or all subtypes using pC1-SS2 led to sleep patterns of mated females. In contrast, simultaneous activation of pC1 subtypes using the same drivers led to virgin-like egg-laying and sexual receptivity behaviors^25,26^. Interestingly, pC1d is the only subtype involved in female aggression^45^. These findings suggest that pC1 subtypes impact distinct sets of female behavior by sending outputs to different downstream circuits.

We found that activating pC1a,c,e or pC1e neurons reduced the calcium levels of dFB-projecting neurons, suggesting that postmating sleep suppression is achieved through dFB inhibition. Numerous studies have shown the critical role of dFB-projecting neurons in sleep regulation under various conditions. Their activation leads to increased sleep by acting as effectors of homeostatic sleep needs^39,46^, and their enhanced activity underlies increased sleep in young flies^53^ and in sickness^54^. They also receive information from neurons involved in sleep suppression due to chronic social isolation^55^, and GABAergic inputs to dFB contribute to temperature-dependent sleep plasticity^56^. These findings suggest that neurons projecting to the dFB integrate inputs from multiple circuits that sense changes in internal states and environmental contexts to adjust sleep. We propose that dFB neurons integrate information about the mating status and sleep drive to balance achieving goal-directed behaviors and avoiding excessive sleep pressure (Figure 6D).

Several neural pathways provide information about food substrate to regulate egg laying, including gustatory^16,57^ and olfactory signaling pathways^58^. A previous study showed that the neuronal activity of OviIN neurons is modulated by the sucrose content and hardness of the egg-laying substrate^25^, suggesting that they are involved in integrating mating status and food composition signals (Figure 6D). Our results show that food composition influences the sleep-suppressive role of pC1a,c,e, and pC1e neurons, suggesting that these or downstream neurons are also involved in integrating information regarding the mating status and food composition. How the food composition information is integrated with mating status and sleep drive signals to regulate postmating sleep is an important topic for future investigation.

Previous studies suggest that sleep changes due to reproductive needs are sexually dimorphic. For example, males lose sleep in the presence of sexual partners in mice, arctic sandpipers, and flies^3,48,50,59^. On the other hand, females exhibit changes in sleep patterns during pregnancy and after the birth of the offspring. Pregnancy-related sleep changes have been observed in humans and rodents^60,61^. Notably, female rats exhibit sleep changes after mating with vasectomized males^62^. Similarly, female flies exhibit postmating sleep suppression after mating with males lacking sperm^33,49^. These results suggest that postmating sleep changes can occur independent of fertilization. Several mammalian species exhibit postpartum sleep reduction^63–65^, and sleep loss due to offspring care is also observed in worker bees^66^. The diversity of the reproductive modulation of sleep may reflect the sex- and species-specific reproductive roles of individual organisms. We have shown that food context modulates the sleep-reproduction balance in mated female flies. It would be interesting to investigate whether similar modulations occur in other species.

## Acknowledgments

We thank the Janelia Fly Facility and the Bloomington Stock Center for fly stocks; Dr. Bill Joiner for the SleepLab software; Dr. Leslie Griffith for hidden Markov model analysis scripts, and members of the Koh lab for helpful discussions. This work was supported in part by a Pew Latin American Fellowship (to J.M.D.), a Japanese Society for Promotion of Science Fellowship (to S.I.), a grant from the National Institutes of Health (R01NS109151 to K.K.), and funds from Thomas Jefferson University Center for Synaptic Biology (to K.K.).

## Author contributions

Conceptualization & Methodology, J.M.D. J.R.B., and K.K. Investigation, Formal Analysis and Visualization, J.M.D., J.R.B., S.I., K.J.K., B.P.J., and D.J.S.A. Writing – Original Draft, J.M.D. and K.K. Writing – Review & Editing, J.R.B., S.I., K.J.K., B.P.J. and D.J.S.A. Funding Acquisition, J.M.D., S.I. and K.K. Supervision and Project Administration, K.K.

## Methods

### Key Resources Table

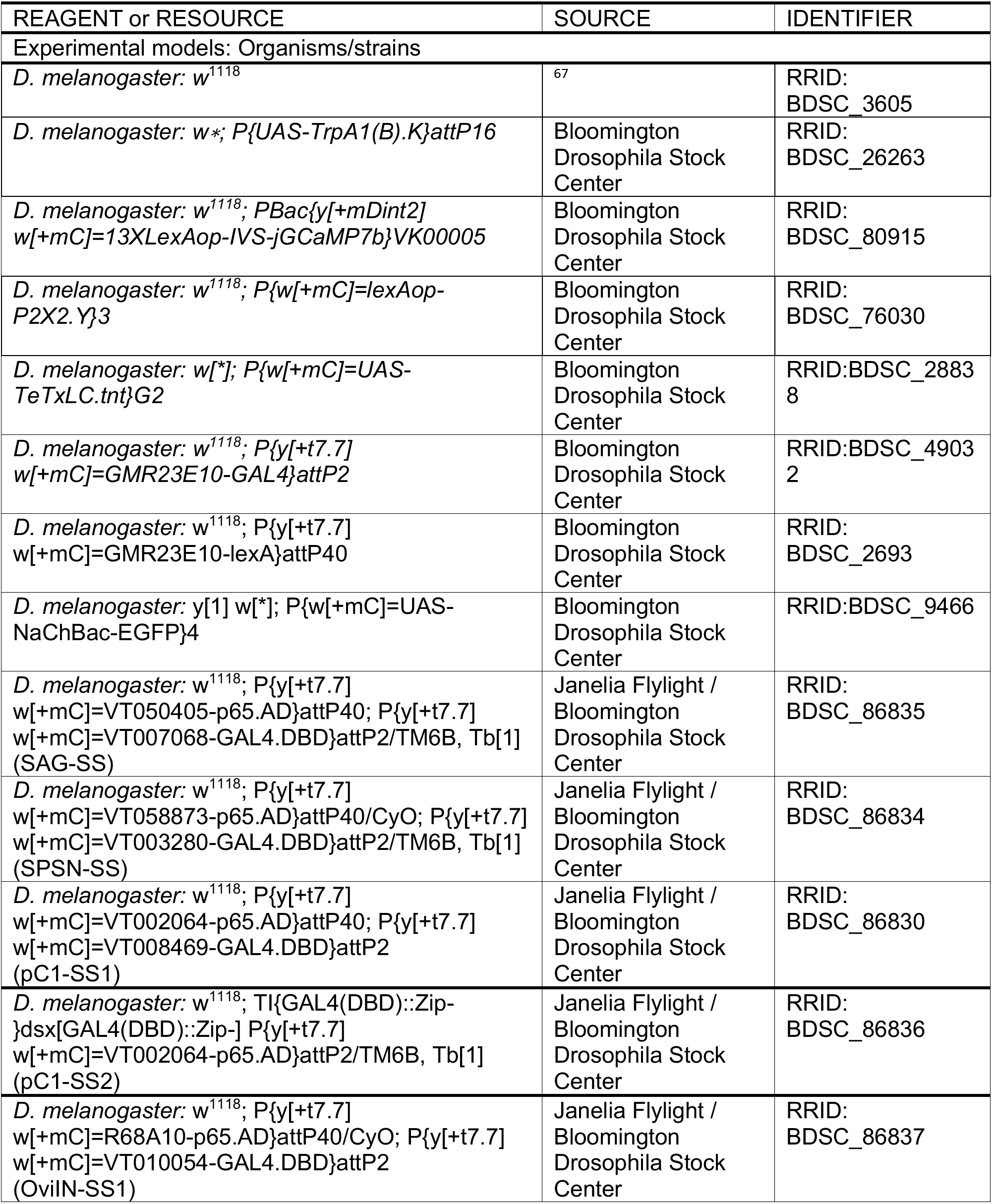
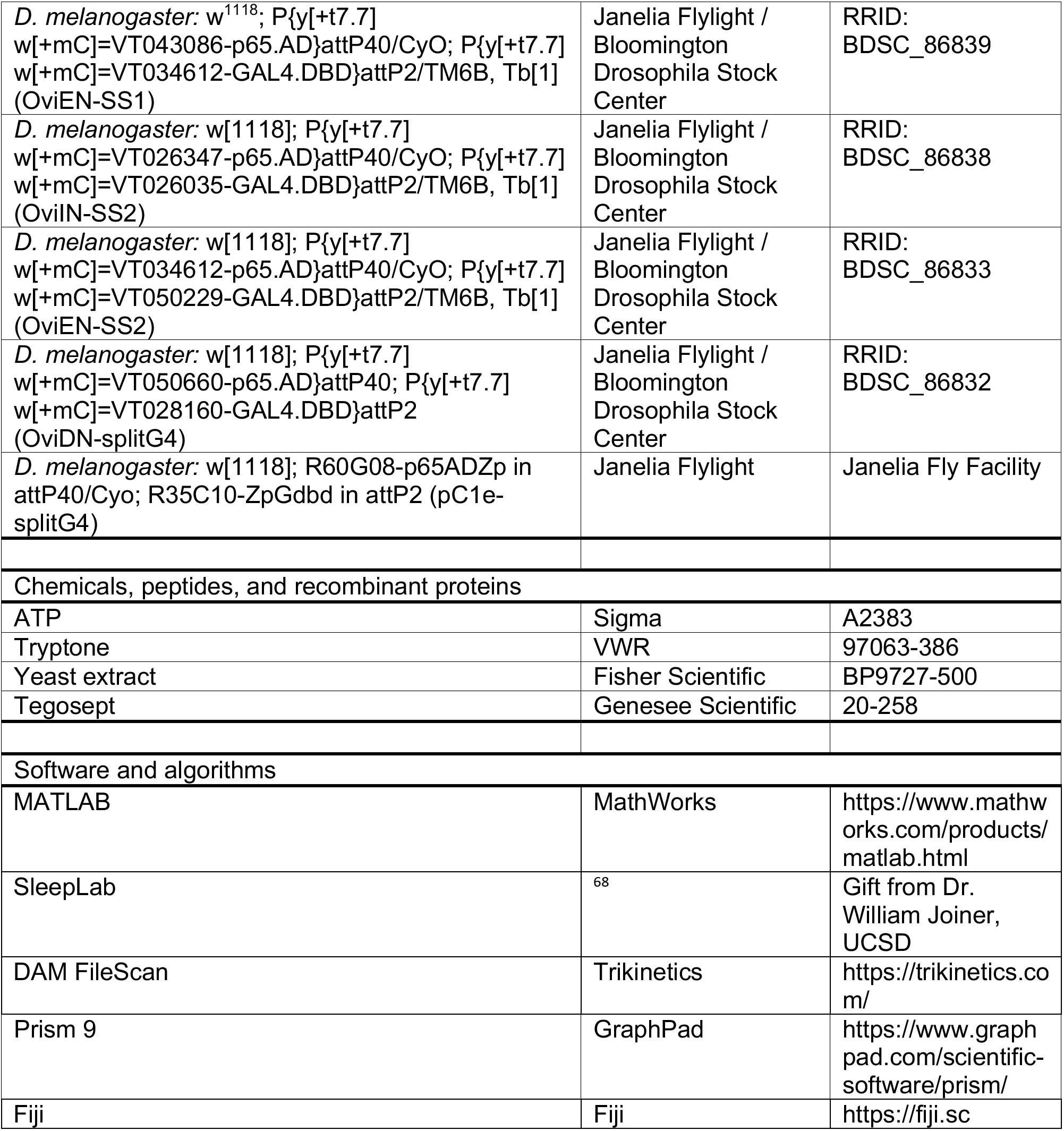

| REAGENT or RESOURCE | SOURCE | IDENTIFIER |
| --- | --- | --- |
| Experimental models: Organisms/strains |  |  |
| *D. melanogaster*: $w^{1118}$ | <sup>67</sup> | RRID:<br>BDSC_3605 |
| *D. melanogaster*: $w^*$; $P\{UAS-TrpA1(B).K\}attP16$ | Bloomington<br>Drosophila Stock<br>Center | RRID:<br>BDSC_26263 |
| *D. melanogaster*: $w^{1118}$; $PBac\{y[+mDint2]$<br>$w[+mC]=13XLexAop-IVS-jGCaMP7b\}VK00005$ | Bloomington<br>Drosophila Stock<br>Center | RRID:<br>BDSC_80915 |
| *D. melanogaster*: $w^{1118}$; $P\{w[+mC]=lexAop-$<br>$P2X2.Y\}3$ | Bloomington<br>Drosophila Stock<br>Center | RRID:<br>BDSC_76030 |
| *D. melanogaster*: $w[*]$; $P\{w[+mC]=UAS-$<br>$TeTxLC.tnt\}G2$ | Bloomington<br>Drosophila Stock<br>Center | RRID:BDSC_2883<br>8 |
| *D. melanogaster*: $w^{1118}$; $P\{y[+t7.7]$<br>$w[+mC]=GMR23E10-GAL4\}attP2$ | Bloomington<br>Drosophila Stock<br>Center | RRID:BDSC_4903<br>2 |
| *D. melanogaster*: $w^{1118}$; $P\{y[+t7.7]$<br>$w[+mC]=GMR23E10-lexA\}attP40$ | Bloomington<br>Drosophila Stock<br>Center | RRID:<br>BDSC_2693 |
| *D. melanogaster*: $y[1] w[*]$; $P\{w[+mC]=UAS-$<br>$NaChBac-EGFP\}4$ | Bloomington<br>Drosophila Stock<br>Center | RRID:BDSC_9466 |
| *D. melanogaster*: $w^{1118}$; $P\{y[+t7.7]$<br>$w[+mC]=VT050405-p65.AD\}attP40$; $P\{y[+t7.7]$<br>$w[+mC]=VT007068-GAL4.DBD\}attP2/TM6B, Tb[1]$<br>(SAG-SS) | Janelia Flylight /<br>Bloomington<br>Drosophila Stock<br>Center | RRID:<br>BDSC_86835 |
| *D. melanogaster*: $w^{1118}$; $P\{y[+t7.7]$<br>$w[+mC]=VT058873-p65.AD\}attP40/CyO$; $P\{y[+t7.7]$<br>$w[+mC]=VT003280-GAL4.DBD\}attP2/TM6B, Tb[1]$<br>(SPSN-SS) | Janelia Flylight /<br>Bloomington<br>Drosophila Stock<br>Center | RRID:<br>BDSC_86834 |
| *D. melanogaster*: $w^{1118}$; $P\{y[+t7.7]$<br>$w[+mC]=VT002064-p65.AD\}attP40$; $P\{y[+t7.7]$<br>$w[+mC]=VT008469-GAL4.DBD\}attP2$<br>(pC1-SS1) | Janelia Flylight /<br>Bloomington<br>Drosophila Stock<br>Center | RRID:<br>BDSC_86830 |
| *D. melanogaster*: $w^{1118}$; $TI\{GAL4(DBD)::Zip-$<br>$\}dsx[GAL4(DBD)::Zip-] P\{y[+t7.7]$<br>$w[+mC]=VT002064-p65.AD\}attP2/TM6B, Tb[1]$<br>(pC1-SS2) | Janelia Flylight /<br>Bloomington<br>Drosophila Stock<br>Center | RRID:<br>BDSC_86836 |
| *D. melanogaster*: $w^{1118}$; $P\{y[+t7.7]$<br>$w[+mC]=R68A10-p65.AD\}attP40/CyO$; $P\{y[+t7.7]$<br>$w[+mC]=VT010054-GAL4.DBD\}attP2$<br>(OviIN-SS1) | Janelia Flylight /<br>Bloomington<br>Drosophila Stock<br>Center | RRID:<br>BDSC_86837 |
| *D. melanogaster*: w<sup>1118</sup>; P{y[+7.7] w[+mC]=VT043086-p65.AD}attP40/CyO; P{y[+7.7] w[+mC]=VT034612-GAL4.DBD}attP2/TM6B, Tb[1] (OviEN-SS1) | Janelia Flylight /<br>Bloomington<br>Drosophila Stock<br>Center | RRID:<br>BDSC_86839 |
| *D. melanogaster*: w[1118]; P{y[+7.7] w[+mC]=VT026347-p65.AD}attP40/CyO; P{y[+7.7] w[+mC]=VT026035-GAL4.DBD}attP2/TM6B, Tb[1] (OviIN-SS2) | Janelia Flylight /<br>Bloomington<br>Drosophila Stock<br>Center | RRID:<br>BDSC_86838 |
| *D. melanogaster*: w[1118]; P{y[+7.7] w[+mC]=VT034612-p65.AD}attP40/CyO; P{y[+7.7] w[+mC]=VT050229-GAL4.DBD}attP2/TM6B, Tb[1] (OviEN-SS2) | Janelia Flylight /<br>Bloomington<br>Drosophila Stock<br>Center | RRID:<br>BDSC_86833 |
| *D. melanogaster*: w[1118]; P{y[+7.7] w[+mC]=VT050660-p65.AD}attP40; P{y[+7.7] w[+mC]=VT028160-GAL4.DBD}attP2 (OviDN-splitG4) | Janelia Flylight /<br>Bloomington<br>Drosophila Stock<br>Center | RRID:<br>BDSC_86832 |
| *D. melanogaster*: w[1118]; R60G08-p65ADZp in attP40/Cyo; R35C10-ZpGdbd in attP2 (pC1e-splitG4) | Janelia Flylight | Janelia Fly Facility |
| Chemicals, peptides, and recombinant proteins |  |  |
| ATP | Sigma | A2383 |
| Tryptone | VWR | 97063-386 |
| Yeast extract | Fisher Scientific | BP9727-500 |
| Tegosept | Genesee Scientific | 20-258 |
| Software and algorithms |  |  |
| MATLAB | MathWorks | https://www.mathworks.com/products/matlab.html |
| SleepLab | <sup>68</sup> | Gift from Dr. William Joiner, UCSD |
| DAM FileScan | Trikinetics | https://trikinetics.com/ |
| Prism 9 | GraphPad | https://www.graphpad.com/scientific-software/prism/ |
| Fiji | Fiji | https://fiji.sc |

### Resource Availability

#### Lead Contact

Further information and requests for resources and reagents should be directed to and will be fulfilled by the Lead Contact, Kyunghee Koh.

#### Material Availability

All unique reagents generated in this study are available from the Lead Contact upon request without restriction.

#### Data and Code Availability

All data supporting the findings of this study are included within the article and Supplementary Information files.

### Experimental Model and Subject Details

#### Fly Husbandry

Flies were raised on standard Drosophila culture media in a 12 h:12 h light:dark (LD) cycle. Standard culture media (referred to as normal food) is composed of 6.56% w/v cornmeal, 2.75% w/v yeast, 0.81% w/v agar, 6.48% v/v molasses, 0.93% v/v propionic acid and 0.25% v/v tegosept (anti-fungal agent, Genesee Scientific, El Cajon, CA). Flies were raised at 25°C except where noted. Virgin females and were collected shortly after eclosion and aged in groups of ~32 for 2-4 days. Before sleep analysis, 16 virgin female flies of all genotypes were grouped with 16 wild-type (iso31) virgin males to generate mated females. Flies in the virgin condition were transferred to new vials at the same time. Flies in all food conditions were raised on normal food. Genotypes used in this study are listed in the Key Resources Table.

### Method Details

#### Sleep Analysis

Flies were monitored in a 12 h:12 h LD cycle at 25°C except when noted. 3- to 5-day old virgin or mated female flies were placed in glass tubes containing one of the following food substrates: normal food, 5% w/v sucrose-2% w/v agar food, and 5% w/v sucrose-2% w/v agar food supplemented with 2.75% w/v yeast extract (Fisher Scientific, Waltham, MA), 2.75% w/v tryptone (VWR, Radnor, PA), or additional 2.75% sucrose. Activity data were collected in 1 min bins using *Drosophila* Activity Monitoring (DAM) System (Trikinetics, Waltham, MA). Single-beam monitors were used except where noted, with the border of the food placed ~0.5 fly body length from the infrared (IR) sensor. Beam breaks from single-beam monitors or intra-beam movements from multi-beam monitors with 17 IR detectors were used for sleep analysis. For experiments involving TrpA1, flies were raised in LD at 22°C and monitored for ~16 h at 22°C to determine baseline levels, followed by 1 day at 27°C or 29°C to activate the TrpA1 channel. For sleep deprivation experiments, flies placed in single-beam monitors were deprived of sleep using mechanical stimulation. A multi-tube vortexer fitted with a mounting plate (Trikinetics, Waltham, MA) was used to apply mechanical stimulation for 3 s every min. To analyze the contribution of larval movement to sleep measurements, we placed mated females in glass tubes with normal food at ~ZT 8 and allowed them to lay eggs for ~24 h. Then adult flies were removed, and beam crossings were recorded from the tubes containing eggs and hatched larvae. Sleep was defined as a period of inactivity lasting at least 5 min. Sleep parameters were analyzed using MATLAB-based software, SleepLab (William Joiner). The hidden Markov model for sleep depth analysis was run in MATLAB as described^34^. Location-based activity heatmaps were constructed from multi-beam DAM data using custom MATLAB scripts.

#### Egg Laying Analysis

Mated females were placed in vials containing either normal food or 5% sucrose in groups of 4 at ~ZT 8 of Day 0. At ZT 0 of Day 1 (which matches the day used for sleep analysis), flies were transferred to new tubes of the same food type, and then transferred again to the same food type at ZT 12 and left until ZT 0 of Day 2. The number of laid eggs in each vial was counted for both daytime (ZT 0-12) and nighttime (ZT 12-24) on Day 1.

#### Calcium Imaging

3- to 5-day-old virgin and mated females entrained to LD cycles were anesthetized on ice and dissected in adult hemolymph-like solution (AHL, 108 mM NaCl, 5 mM KCl, 2 mM CaCl2, 8.2 mM MgCl2, 4 mM NaHCO3, 1 mM NaH2PO4, 5 mM trehalose, 10 mM sucrose, 5 mM HEPES, pH 7.5, 265 mOsm)^69^. Dissected brains were mounted on a glass-bottom chamber containing AHL solution. The flow rate of the solutions was controlled by a custom-built gravity-dependent perfusion system. Images were acquired on a Leica SP8 confocal microscope every 5 s for 4.5 min. 2.5 mM ATP in AHL was delivered for 1 min after 0.5 min of baseline measurements. Average intensity projections were computed in FIJI, and the fluorescence intensity of the cell body area was quantified. The mean intensity during the 30 s period prior to ATP perfusion was used as the baseline measurement, F_0_. Normalized ΔF, (F-F_0_/F_0_, was computed for each time point.

#### Quantification and Statistical Analysis

Statistical tests were performed using Prism 9 (GraphPad, San Diego, CA). For experiments involving two factors, two-way ANOVAs were performed to test for the main and interaction effects, followed by post hoc tests to compare specific pairs of groups. Dunnett’s post hoc tests were used when comparing flies carrying both a driver and a USA transgene against parental controls carrying the driver alone or the transgene alone. Tukey’s or Sidak’s post hoc tests were used in all other cases. Student’s t test was employed to compare pairs of groups. Details of statistical tests, including p values and n, can be found in figure legends.

